# Structural covariation between cerebellum and cerebral cortex is atypically modulated by thalamus in autism spectrum disorder

**DOI:** 10.1101/2023.10.16.562588

**Authors:** Federico d’Oleire Uquillas, Esra Sefik, Bing Li, Matthew A. Trotter, Kara Steele, Jakob Seidlitz, Rowen Gesue, Mariam Latif, Tristano Fasulo, Veronica Zhang, Mikhail Kislin, Jessica L. Verpeut, Jonathan D. Cohen, Jorge Sepulcre, Samuel S.-H. Wang, Jesse Gomez

## Abstract

Despite its strong interconnectivity with the cerebral cortex, the influence of the human cerebellum on neocortical structure and its role in the development of neuropsychiatric disorders is unclear. Because cerebellar damage in early postnatal life creates a high risk for autism spectrum disorder (ASD), we investigated inter-relationships in cerebello-cerebral morphology. Leveraging a large structural brain MRI dataset in neurotypical children (n=375) and those diagnosed with ASD (n=373), we examined whether volumetric variation in cerebellar structure across individuals was correlated with neocortical variation during development, modeling the thalamus as a moderating coupling factor. We found negative covariation between cerebellar cortical regions and thalamic/sensorimotor neocortical regions, and positive covariation between thalamic and sensorimotor neocortical regions. This pattern aligned with the major disynaptic path of cerebellar inhibition to thalamocortical excitation. Examining the dependence of this structural covariation on ASD diagnosis, we found that neurotypical and ASD children displayed inverted hemispheric biases. In ASD, the thalamus moderated structural associations between the left cerebellum and right sensorimotor cortex. For neurotypical children, right cerebellum and left sensorimotor cortex were coupled. Notably, structural coupling between cerebellum, thalamus, and neocortex was strongest in younger childhood and waned by early adolescence, a time during which behavioral differences were smallest between typically developing and autistic children. In addition to the sensorimotor cortex, cerebellar dentate nuclei in ASD displayed greater coupling to broad neocortical regions, and these nuclei related to cognitive function differently such that greater dentate nuclear volume was associated with greater behavioral impairment in ASD but not in controls. Graph analyses demonstrated that the cerebello-thalamocortical network was more densely and less prolifically interconnected in ASD than in typical development. Taken as a whole, our study reveals a developmental interplay between the cerebellum, thalamus, and neocortex which differs in ASD from neurotypical children. This pattern is consistent with a model of ASD in which early developmental influences of cerebellar output on brain maturation are specifically moderated by cerebello-thalamocortical pathways.

## Introduction

The cerebellum plays an important role in a wide array of cognitive functions. A recent surge in work in cognitive neuroscience supports a long history of clinical findings demonstrating that adult cerebellar damage can lead to cognitive and affective dysfunction^1^. Moreover, perinatal cerebellar damage presents the largest non-heritable risk ratio for a later diagnosis of autism spectrum disorder (ASD)^2^. Perinatal cerebellar hemorrhage events appear to be overrepresented in ASD and are often found in premature and low birth weight infants^3–6^. These and related findings have led to the developmental diaschisis hypothesis^2^, which proposes that disruption of cerebellar function during sensitive periods of brain development result in long-distance changes in neocortical maturation.

Evidence for the potential of long-distance influence of cerebellar injury originates from observations of diaschisis in adults, defined as the impairment or dysfunction of brain regions distant from a site of primary injury. Cerebellar hemorrhagic events or stroke have been associated with reduced hypometabolism in the contralateral cerebral hemisphere^7–11^. The importance of cerebello-cortical pathways is also evident in patients with unilateral cerebellar lesions, who show reduced excitability in the form of higher magnetic stimulation thresholds required for motor stimulation of contralateral motor cortex^12^. A different imaging study in adult patients who were treated for cerebellar tumors in their childhood found that increased prefrontal cortex gray matter volume was related to decreased performance on processing speed^13^, indicative of cerebellar influence on both cerebral structure and function. Influences are reciprocal, as crossed cerebellar diaschisis is observed in response to supratentorial hemispheric atrophy^14,15^. However, while interconnectivity between the neocortex and cerebellum underlies a number of medical pathologies, whether and how it is altered during neurodevelopment in children diagnosed with ASD without obvious cerebellar injury is unclear.

The cerebellum influences the rest of the brain via Purkinje cell axons, which project from cerebellar cortex to inhibit the deep cerebellar nuclei and other structures. Of these output structures, the dentate nuclei are of specific interest because they are the largest of the deep nuclei in humans and because they are more susceptible to insult than the other nuclei^16^. The deep nuclei send excitation to the thalamus, the principal gateway to the cerebral cortex^17–19^. Viral transsynaptic tract-tracing studies in mice have shown that the cerebello-thalamic paths project to somatomotor, somatosensory, and association cortex^20^. In addition, atypical functional connectivity and sensory response alterations in ASD have been reported in the sensorimotor cortex (SMC)^21^, one of the largest projection zones of cerebellar output^22^.

Taken together, these findings suggest the possibility that the cerebellum’s influence in ASD can be investigated by examining the joint variation in the structure or function of cerebellum, thalamus, and cerebral cortex. One powerful tool for conducting such an investigation is structural covariance^23^. Pediatric functional neuroimaging is difficult to collect reliably in sensitive populations like those with ASD. Anatomical quantification of the volumes of brain structures provides a robust measure. Volumes of interconnected brain regions covary across individuals and are thought to reflect the cumulative effect of coordinated growth and pruning processes. Using anatomical correlations in this way is supported by the observation that nodes of functional networks, such as the default mode network (DMN), dynamically co-vary in their structure during brain maturation^23^. Moreover, dynamic shifts occur in the network topology of cognitive cerebral networks from childhood to adolescence^24^. Would atypical structural connectivity across an extended cerebello-cerebral network in ASD potentially disrupt the structural covariation in volume of these structures? Answering such a question requires a large dataset of both ASD and neurotypical children, in turn requiring automated methods for segmenting cerebellum, neocortex, and the thalamus into their constituent regions and nuclei.

To meet this challenge, we used recent neuroanatomical parcellation tools to analyze one of the largest collections of structural brain images of ASD patients to date, obtained from over 700 individuals typically developing (TD) and diagnosed with ASD. We controlled for sex, age, and cranial size to test whether remaining variation in cerebellar and neocortical volumes showed ASD-dependent covariation. We further tested whether the thalamus acted as a moderator of cerebellar-neocortical co-variation across development, whether cerebellar structure could account for behavioral differences in ASD, and whether cerebellar output influences large-scale network properties.

## Materials and methods

### Participants

This study considered a total of 1050 participants from the publicly available ABIDE-I and ABIDE-II dataset repositories^25,26^. Inclusion in this study was restricted to participants up to age 19, above which available ASD cases decreased in number. Individual T1-weighted scans deemed unusable due to poor spatial resolution or image quality were excluded. The final cohort consisted of n=373 individuals with an ASD diagnosis as determined by a trained clinician (64 female, 309 male, age range: 7 – 19 years, sd age: 2.80, mean age: 12.59, median age: 12.20), and n=375 typically developing (TD) individuals (112 female, 263 male, age range: 7 – 19 years, sd age: 2.80, mean age: 12.05, median age: 11.56).

### Clinical Assessment

Criteria for assessing diagnosis and symptom severity included the Autism Diagnostic Interview-Revised (ADI-R)^27,28^ and the Autism Diagnostic Observation Schedule (ADOS)^29,30^. Original inclusion criteria used Diagnostic and Statistical Manual of Mental Disorders (DSM-5) codes in addition to DSM-IV criteria for autism for those enrolled after 2013. We included TD subjects based on absence of any current Axis-I disorders in children as per KSADS-PL^31^ or DSM-IV-TR SCID for adults^32^, group-matched to ASD on age. Chronic systemic medical or psychiatric conditions or MRI contraindications were grounds for exclusion.

### Cognitive Assessment

The Behavior Rating Inventory of Executive Function (BRIEF)^33^ was available to measure impairment across a variety of cognitive domains (Inhibition, Shifting, Working Memory, Organization, Emotional Regulation, Initiation, and Planning subscales), with a total score across all domains in addition to individual subsection scores, for each participant.

### Magnetic Resonance Imaging

T1-weighted imaging data was acquired within 3 months of diagnostic assessment using a 3D-MPRAGE sequence. Scan parameters can be publicly found at the ABIDE source site: http://fcon_1000.projects.nitrc.org/indi/abide/scan_params/. We processed T1-weighted cerebral^34,35^ and thalamic nuclear data^36^ using FreeSurfer 7.1.1 (http://surfer.nmr.mgh.harvard.edu/), and used the Human Connectome Project multi-modal parcellation atlas (HCP-MMP^37^) to produce regional estimates of neocortical gray matter for use in statistical analyses, defined from labels of the multi-modal parcellation in each individual’s brain.

We processed each individual’s cerebellum using the spatially unbiased atlas of the cerebellum and brainstem (SUIT) toolbox^38^ for SPM12 (http://www.fil.ion.ucl.ak.uk/spm/). For deep cerebellar nuclei, we focused on the dentate nucleus^39^, the largest of the three major nuclei. Because local inconsistencies in the identification of anatomical structures can add variation to the structural covariance analysis, trained raters visually inspected each parcellation. Large errors that could not be corrected by trained anatomists led to exclusion from the study. Of the remaining participant data, minor labeling inaccuracies were manually corrected to ensure cerebellar volume estimates were accurate.

### Statistical Analysis

#### Sample Characteristics

We quantified associations between sample characteristics using R (v.4.0.4; https://www.R-project.org) (**Table 1**). Normality for brain-derived regional volumes was estimated using the two-sample Kolmogorov-Smirnov test, which compares two samples to determine if the variable in question is normally distributed, and if the distributions between two samples are different from one another. Measures including *estimated total intracranial volume (eTIV)*, *total cerebral gray matter volume*, and *total cerebellar gray matter volume* were normally distributed and did not differ from one another (Kolmogorov-Smirnov, *p*>0.1), allowing the use of subsequent tests involving Pearson correlations and general linear models.

**Table 1.**

|  | *Autism Cohort* | *Neurotypical Cohort* | *p-value* |
| --- | --- | --- | --- |
| n | 373 | 375 |  |
| Age | 12.59 (2.80) | 12.05 (2.80) | 0.009 |
| Sex No. (%) = F | 64 (17.2) | 112 (29.9) | < 0.001 |
| BRIEF: Composite Score | 63.62 (9.32) | 47.01 (6.44) | < 0.001 |
| VIQ | 106.04 (18.60) | 112.90 (14.97) | < 0.001 |
| eTIV | 1,573,797.36 (199797.96) | 1,536,434.64 (187725.49) | 0.009 |
| Cerebellar Cortex Volume [mm<sup>3</sup>] | 8,466.59 (450.24) | 8,536.76 (446.49) | 0.047<sup>a</sup> |
| Dentate Cerebellar Nuclei Volume [mm<sup>3</sup>] | 1,963.05 (158.41) | 1,953.67 (158.14) | 0.435<sup>a</sup> |
| Thalamic Nuclei Volume [mm<sup>3</sup>] | 31.16 (6.32) | 30.43 (5.62) | 0.546<sup>a</sup> |
| Sensorimotor Cortex Volume [mm<sup>3</sup>] | 291,445.21 (88,554,948.06) | 269,648.00 (74,348,103.34) | 0.0003<sup>a</sup> |

Note: Data is reported as mean (standard deviation). Abbreviations: eTIV, estimated total intracranial volume; VIQ, verbal IQ ability as measured by the American National Adult Reading Test. <sup>a</sup>Model corrected for sex, age, and eTIV.

#### Structural Covariance Analysis

To map volumetric associations between cerebral, cerebellar, and thalamic ROIs, we calculated Pearson correlations across all participants in a group between every region pair to obtain a 78×78 correlation matrix. For neocortex, we used a sensorimotor cortex (SMC) meta-ROI that encapsulated primary motor and areas 1L, 2L, and 3aL of sensory cortex. For the cerebellum, we used cerebellar lobules and dentate nuclei derived from SUIT (26 regions). For the thalamus, we used 25 thalamic nuclei per hemisphere (50 regions).

#### Cerebellar Nuclei x Sensorimotor Thalamic Nuclei Interaction Analysis

We performed moderation analysis to further understand how sensorimotor thalamic nuclei affected the relationship between dentate nuclei and distal SMC, in the form of a three-way interaction (SMC ~ DN x thalamic nuclei x diagnosis). Demographic variables for age, eTIV, and sex assigned at birth were included as covariates in all linear regression models. Homoscedasticity assumptions were considered, and robust standard error estimates were calculated. Standardized β-coefficients are reported for all models, along with bias-corrected and accelerated (BCa) bootstrapped 95% confidence intervals, and Cohen’s *f* effect sizes for partial effects in multivariate regressions.

#### Probing Thalamic Interactions Across Development

We split the study sample by diagnosis to examine the thalamic moderation effect across the observed developmental window (SMC ~ DN x thalamic nuclei x age, separately within each group). Rather than assume developmental processes were linear, we compared linear, quadratic and cubic models using analysis of variance and Akaike Information Criterion (AIC) to identify a parsimonious best-fit model.

#### Cognitive Trajectories Analysis and Anatomical Correlates

To characterize cognitive trajectories we analyzed a subset of individuals with complete cognitive data (n=178: 47 female, 131 male, age range: 7 – 18 years, mean age: 11.3, sd age: 2.5, median age: 10.9) for associations between age and performance on the BRIEF battery. The battery included subscores measuring inhibition, working memory, shifting, organization, and planning scores. We also considered a participant’s overall performance on the battery through an average of all subscales. BRIEF ratings reflect T-scores (population mean from normative samples, M = 50; SD = 10), with scores ≥ 70 considered to be clinically significant. We compared linear, quadratic and cubic age fits to identify the best fit as defined by analysis of variance and AIC. To test whether BRIEF cognitive scores correlated with ROI anatomy along the ascending cerebello-thalamo-SMC pathway in an ASD-dependent manner, a post-hoc model was used (BRIEF composite ~ sex + age + eTIV + DN x Group) with dentate nuclear volume as predictor of cognition.

#### Exploratory Cerebral Associations with Cerebellum Analysis

To better understand how integrity of the cerebellar dentate nuclei related to cerebral regions other than our *a priori* hypothesis involving the SMC, we conducted a post-hoc vertex-wise cortical thickness analysis. False-positive rates in our surface-based anatomical analysis were controlled (*p*-corrected<0.05) using 1000 permutation-based simulations^40^. To further compare overall dentate nuclei-covariance network properties such as density, hubness and graph degree, we constructed an average covariance unthresholded network for each cohort, using volumes from the 180-neocortical Glasser Atlas regions, thalamic nuclei, and cerebellar regions, such that each node in the network represented each region, and every edge represented covariance between two nodes. Networks were visualized using force-directed Kamada-Kawai plots, which simulate a physical system where nodes are like charged particles and edges act like springs, placing nodes with high covariance inward, while placing nodes with low covariance to the rest of the network around the periphery.

### Code Availability

Code written to conduct these analyses are available on GitHub. Raw data scans are publicly available (http://fcon_1000.projects.nitrc.org/indi/abide/).

## RESULTS

From each participant, we sent the T1-weighted neuroanatomical image through three parallel processing pipelines, one to identify and parcellate the cortical ribbon^34,35^, one to segment the thalamus into its constituent nuclei^36^, and another to parcellate the cerebellum^38^, including the identification of its dentate nuclei^39^. This resulted in a final dataset of *n=375* TD and *n=373* ASD participants.

In line with the rate of ASD in males being 3 to 4 times greater than in females, the distribution of sex in the ASD cohort differed significantly from that in the TD group, as per chi-squared test (*p*<0.001). Groups also differed in their average age (TD mean age: 12.20, ASD mean age: 12.05; β=−0.61, se=0.22, Cohen’s *f*=0.11, *p*=0.006). Controlling for sex and age, groups did not differ in their eTIV (*p*=0.25).

### Structural Covariance Between Cerebellum, Thalamus, and Sensorimotor Cortex

Covariance analysis can identify whether nodes of a network act as if they have a trophic history with one another (*i.e.*, an increase in the size of one region would correspond to an increase in size of another region). Given that the connectivity between cerebellar nuclei, the thalamus, and the neocortex involve both excitatory and inhibitory connections, both positive and negative covariation might occur between regions.

We first focused on a well-understood circuit in the typically developing cohort between the cerebellum, thalamus, and sensorimotor cortex (SMC), an ascending pathway responsible for integrating sensory information and motor action representations (**Fig 1a**). In this circuit, cerebellar Purkinje cells, a unique class of GABAergic neurons and the only output of the cerebellar cortex, send inhibitory signals to the deep cerebellar nuclei. In our covariance model controlling for sex, age, and eTIV, we also found a negative relationship, where a quadratic term of cerebellar cortex volume best explained dentate nuclear volume values (β=879.14, se=151.2, Cohen’s *f*=0.31, *p*<0.0001) (**Fig 1b**). We focused on the dentate nuclei given previous reports of its abnormal intrinsic functional connectivity with cerebral cortex, including sensorimotor regions, in ASD^41^.

**Figure 1:**
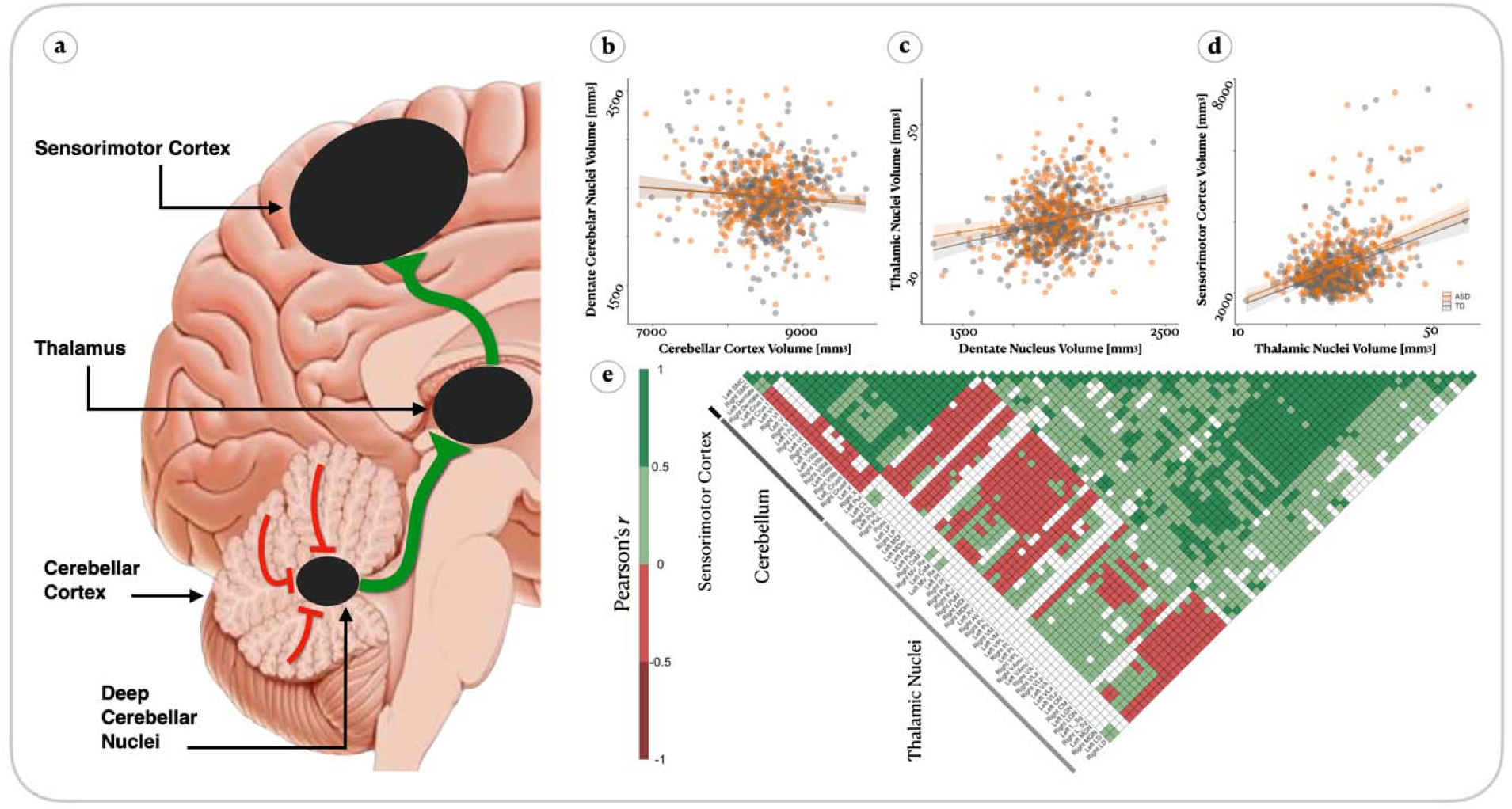
Structural covariation parallels cerebello-thalamo-cortico-pontine circuit. (**a**) Circuit diagram of the cerebellar ascending pathway demonstrating inhibitory influence from cerebellar cortex on deep cerebellar nuclei which synapse onto thalamic nuclei via excitatory projections, ultimately innervating cerebral cortex, (**b-d**) Marginal effects plots of the relationship between cerebellar cortex and dentate cerebellar nuclei (**b**); dentate cerebellar nuclei and thalamic nuclei (**c**); and thalamic nuclei and sensorimotor cortex (**d**), corrected for sex, age, and estimated intracranial volume. (**e**) Correlation matrix, thresholded at *p*<0.0001, in typically developing youth generally showing negative correlations between gray matter volume in cerebellar cortex and deep cerebellar dentate nuclei, positive correlations between dentate nuclei and thalamic nuclei, and positive correlations between thalamic nuclei (central lateral, CL, laterodorsal, LD, medial ventral, MV; combined into a meta-Thalamus ROI) and sensorimotor cortex.

Dentate nuclei send excitatory signals to the thalamus. Likewise, our model showed positive covariation between volumes of dentate nuclei and contralateral sensorimotor thalamic nuclei (β=32.18, se=5.2, Cohen’s *f*=0.32, *p*<0.0001: **Fig 1c**). From thalamus, signals travel via excitatory monosynaptic connections to the neocortex. We found that thalamic and ipsilateral SMC volumes were similarly positively correlated (β=563,400, se=70,990, Cohen’s *f*=0.41, *p*<0.0001: **Fig 1d**). Together, the negative correlation between cerebellar cortex and dentate nuclei and the positive correlations between dentate nuclei, thalamus, and neocortex matched the sign of monosynaptic projections in the ascending pathway. In an unadjusted correlation matrix (*p*<0.0001) between all pairs of regions across SMC, thalamic nuclei, and cerebellar nuclei, the signs of different relationships clustered according to brain structure (**Fig 1e**). These results are consistent with the idea that net excitatory drive from cerebellar nuclei plays a positive role in neocortical growth and/or survival, and that inhibitory output of the cerebellar cortex has an inverse influence.

### Sensorimotor Thalamic Nuclei Moderate Cerebellar-Neocortical Structural Covariance

We next sought to test whether the thalamus moderated the observed covariation between cerebellar and neocortical anatomy in TD. We focused on the sensorimotor circuit leading to SMC. We found that left cerebellar cortex related to right SMC (β=−0.001 se=0.0003, Cohen’s *f*=0.21, *p*<0.0001) [**Table S1a**], and right cerebellar cortex related to left SMC (β=−0.001, se=0.0003, Cohen’s *f*=0.23, *p*<0.0001) [**Table S1b**]. To reduce the number of models, we focused on dentate nuclei-related and sensorimotor cortex-related thalamic nuclei (thalamic nuclei: VPL, VM, VLp, LD, LP) when assessing how thalamus moderated the DN-SMC associations.

Including *group* as a factor in the models, we next tested if the thalamus’ role as a moderator varied by clinical diagnosis. Formally testing for group differences in three-way interaction models (SMC ~ DN x sensorimotor thalamic nuclei x group; one model per hemisphere) controlling for sex and eTIV, we found that the moderating role of the right thalamus among autistic children differed from neurotypical development: lower values of right thalamic nuclei size related to weaker left DN-to right SMC-coupling in ASD (Cohen’s *f*=0.06, *p*=0.024), with no differences between groups in thalamic moderation for the contralateral model (*p*=0.9) (**Table S2**).

To test whether the thalamic moderation effect varied across the maturational process from childhood into adolescence, we examined the thalamic moderation models as three-way interactions with age (SMC ~ DN x thalamic nuclei x age, separately for neurotypical and ASD groups), again controlling for sex and eTIV. These models revealed that greater thalamic size was associated with a stronger coupling between dentate nuclei and sensorimotor cortex size that while observed both in ASD and TD groups (**Fig 2a-b**), were notably different. In the TD group, larger left hemisphere thalamic nuclei were associated with a steeper positive-slope relationship between right dentate nuclei and left sensorimotor cortex volume (95% CI=[−0.05, −0.004], Cohen’s *f*=0.13, *p*=0.0001) (**Table 2**; **Fig 2a**). In the ASD group, the effect was reversed across hemispheres (**Table 2**; **Fig 2b**), such that larger right hemisphere thalamic nuclei were associated with steeper slopes of volumetric coupling between left dentate nuclei and right sensorimotor cortex volume (95% CI=[−0.07, −0.02], Cohen’s *f*=0.15, *p*=0.0002). For ASD children, the thalamic moderation effect was not significant in the left hemisphere after bootstrapping of coefficients (95% CI = −0.05, 0.007). In both groups, the thalamus’ effect on cerebello-neocortical coupling was strongest at the youngest ages, suggesting a prominent role for thalamic influence in early childhood (**Figure 2a-b**). To quantify the age-dependence of thalamic moderation in the TD circuit, we performed a Johnson-Neyman analysis examining at which levels of the moderator the effect of a predictor becomes significant. That is, this analysis evaluated how strongly the thalamus’ moderation effect on cerebellum-SMC coupling differed across the range of ages, effectively asking if the moderation’s significance is constant across development. We found that the slopes of the three-way interaction models with age waned as children approached age 15.

**Figure 2.**
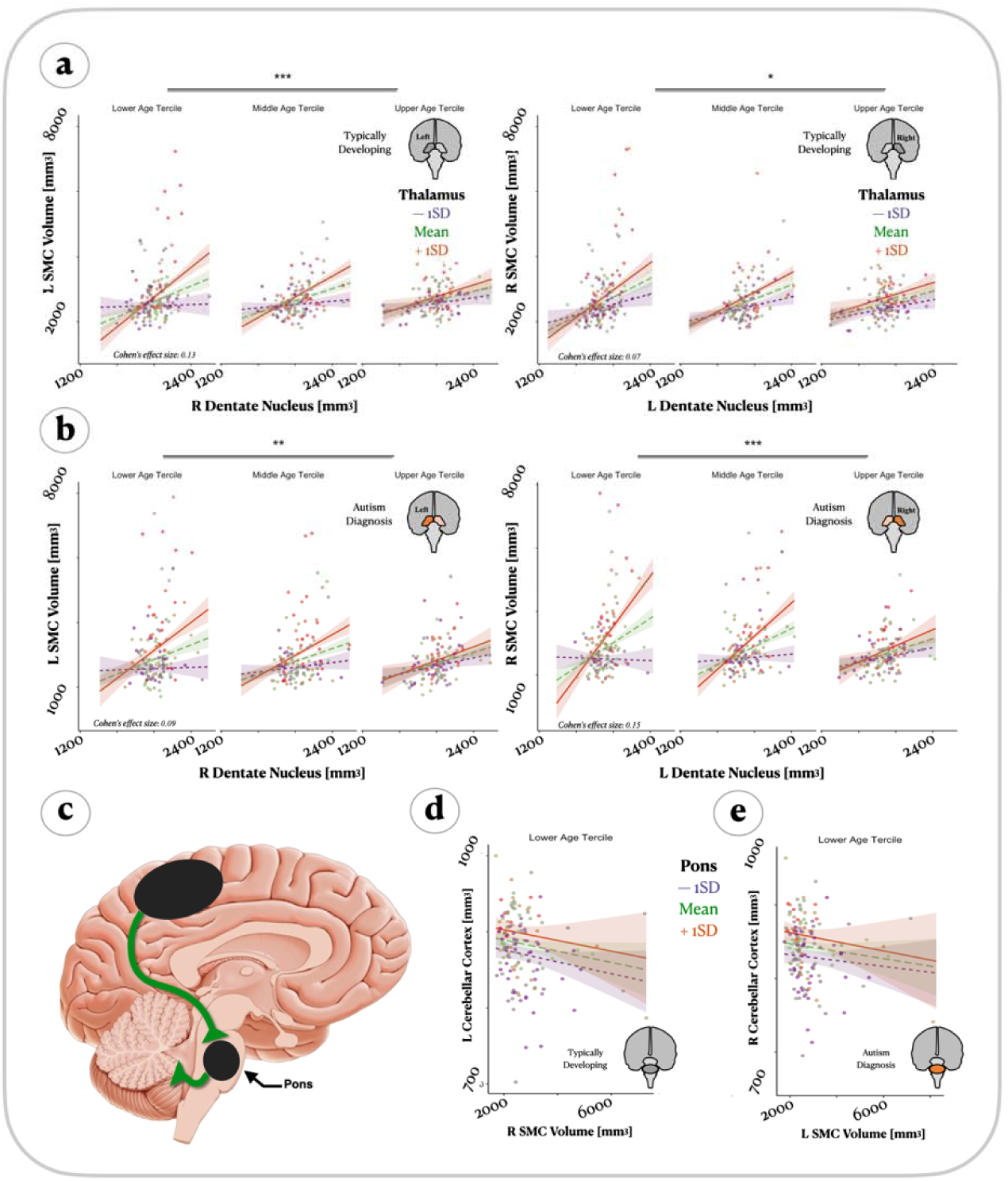
Cerebellar influence on sensorimotor cortex via thalamus and not pons. (**a**) Marginal effects plots, corrected for sex and intracranial volume, of the thalamic nuclei moderation effect on the relationship between gray matter volume in deep cerebellar dentate nuclei and sensorimotor cortex in typically developing children. Model shown per age tercile. (**b**) Similar to panel a, but depicting autism cohort data. The thalamic moderation effect across age in ASD was stronger in the left cerebellar dentate x right thalamus x age model, relative to the contralateral side. In the typically developing cohort, the thalamic moderation effect across age was stronger in the right cerebellar dentate x left thalamus x age model than the contralateral side. Terciles approximated for lower: 9.64 years; middle: 13.74 years; upper: 17.84 years, (**c**) Illustration of the descending pathway between cortex, the pons, and the cerebellum. (**d**) Similar to a but using the pons as a moderator between right somatosensory cortex and left cerebellar cortex. Only lower age tercile shown, other tercile data are similar, (**e**) Similar to **d** but for the other hemisphere: left somatosensory cortex and right cerebellar cortex. Only lower age tercile shown, other tercile data are similar. L, left; R; right.

**Table 2a.**

|  | *Dependent variable:* |  |
| --- | --- | --- |
|  | **Left Sensorimotor Cortex** |  |
|  | *Autism Cohort* | *Controls Cohort* |
| Constant | (5,958.085, 32,645.550)<br>t = 2.835<br>p = 0.005** | (12,825.570, 32,912.030)<br>t = 4.463<br>p = 0.00001*** |
| Right DN | (-16.068, -3.169)<br>t = -2.923<br>p = 0.004**<br>Cohen's *f* = 0.21 | (-15.632, -5.736)<br>t = -4.232<br>p = 0.00003***<br>Cohen's *f* = 0.23 |
| Left Thalamus | (-1,065.479, -190.511)<br>t = -2.813<br>p = 0.005**<br>Cohen's *f* = 0.24 | (-1,122.080, -467.026)<br>t = -4.755<br>p = 0.00001***<br>Cohen's *f* = 0.19 |
| Age | (-2,048.220, -126.643)<br>t = -2.218<br>p = 0.027*<br>Cohen's *f* = 0.05 | (-2,094.690, -587.069)<br>t = -3.486<br>p = 0.0005***<br>Cohen's *f* = 0.02 |
| Biological Sex | (-246.595, 79.819)<br>t = -1.001<br>p = 0.317<br>Cohen's *f* = 0.04 | (-175.676, 40.203)<br>t = -1.230<br>p = 0.219<br>Cohen's *f* = 0.05 |
| eTIV | (-0.0003, 0.0004)<br>t = 0.381<br>p = 0.704 | (-0.0002, 0.0004)<br>t = 0.796<br>p = 0.427 |
| | Cohen's $f = 0.02$ | Cohen's $f = 0.06$ |
| Right DN x Left Thalamus | (0.148, 0.572) | (0.252, 0.571) |
| | $t = 3.333$ | $t = 5.055$ |
| | $p = 0.001^{***}$ | $p = 0.00000^{***}$ |
| | Cohen's $f = 0.13$ | Cohen's $f = 0.15$ |
| Right DN x Age | (0.146, 1.084) | (0.311, 1.062) |
| | $t = 2.568$ | $t = 3.583$ |
| | $p = 0.011^*$ | $p = 0.0004^{***}$ |
| | Cohen's $f = 0.01$ | Cohen's $f = 0.04$ |
| Left Thalamus x Age | (5.153, 69.826) | (24.225, 75.010) |
| | $t = 2.272$ | $t = 3.830$ |
| | $p = 0.024^*$ | $p = 0.0002^{***}$ |
| | Cohen's $f = 0.13$ | Cohen's $f = 0.04$ |
| Right DN x Left Thalamus x Age | (-0.037, -0.006) | (-0.038, -0.013) |
| | $t = -2.652$ | $t = -3.982$ |
| | $p = 0.009^{**}$ | $p = 0.0001^{***}$ |
| | Cohen's $f = 0.09$ | Cohen's $f = 0.13$ |
|  | bca 95% ci = [-0.05, 0.007] | bca 95% ci = [-0.05, -0.004] |
| Observations | 373 | 375 |
| Residual Std. Error | 557.819 (df = 363) | 454.616 (df = 365) |
| *Note:* *$p < 0.05$, **$p < 0.01$, ***$p < 0.001$ | | |

**Table 2b.**

| *Dependent variable:* |  |  |
| --- | --- | --- |
|  | **Right Sensorimotor Cortex** |  |
|  | *Autism Cohort* | *Controls Cohort* |
| Constant | (18,948.370, 52,383.710) | (-252.626, 21,631.260) |
|  | t = 4.181 | t = 1.915 |
|  | p = 0.00003*** | p = 0.056 |
| Left DN | (-27.422, -10.070) | (-10.286, 1.117) |
|  | t = -4.235 | t = -1.576 |
|  | p = 0.00003*** | p = 0.116 |
|  | Cohen's *f* = 0.27 | Cohen's *f* = 0.28 |
| Right Thalamus | (-1,846.548, -704.236) | (-798.041, -78.301) |
|  | t = -4.377 | t = -2.386 |
|  | p = 0.00002*** | p = 0.018* |
|  | Cohen's *f* = 0.27 | Cohen's *f* = 0.26 |
| Age | (-3,323.311, -882.447) | (-1,364.181, 197.904) |
|  | t = -3.377 | t = -1.463 |
|  | p = 0.001*** | p = 0.144 |
|  | Cohen's *f* = 0.08 | Cohen's *f* = 0.01 |
| Biological Sex | (-160.224, 168.951) | (-198.851, 24.710) |
|  | t = 0.052 | t = -1.527 |
|  | p = 0.959 | p = 0.127 |
|  | Cohen's *f* = 0.004 | Cohen's *f* = 0.06 |
| eTIV | (-0.0003, 0.0003) | (-0.0003, 0.0003) |
|  | t = 0.153 | t = 0.125 |
| | $p = 0.879$<br>Cohen's $f = 0.02$ | $p = 0.901$<br>Cohen's $f = 0.02$ |
| Left DN x Right Thalamus | (0.426, 1.017)<br>$t = 4.784$<br>$p = 0.00001^{***}$<br>Cohen's $f = 0.19$ | (0.056, 0.428)<br>$t = 2.552$<br>$p = 0.011^*$<br>Cohen's $f = 0.09$ |
| Left DN x Age | (0.543, 1.814)<br>$t = 3.634$<br>$p = 0.0003^{***}$<br>Cohen's $f = 0.04$ | (-0.117, 0.709)<br>$t = 1.405$<br>$p = 0.160$<br>Cohen's $f = 0.07$ |
| Right Thalamus x Age | (35.375, 120.242)<br>$t = 3.594$<br>$p = 0.0004^{***}$<br>Cohen's $f = 0.13$ | (1.103, 54.158)<br>$t = 2.041$<br>$p = 0.042^*$<br>Cohen's $f = 0.01$ |
| Left DN x Right Thalamus x Age | (-0.066, -0.022)<br>$t = -3.886$<br>$p = 0.0002^{***}$<br>Cohen's $f = 0.15$<br>bca 95% ci = [-0.07, -0.02] | (-0.028, -0.0005)<br>$t = -2.027$<br>$p = 0.043^*$<br>Cohen's $f = 0.07$<br>bca 95% ci = [-0.03, 0.007] |
| Observations | 373 | 375 |
| Residual Std. Error | 533.160 (df = 363) | 456.572 (df = 365) |

Note: \* $p < 0.05$ , \*\* $p < 0.01$ , \*\*\* $p < 0.001$

The thalamus appears to moderate the coupling between cerebellum and cortex differently in ASD and TD. However, is this atypical moderation specific to the thalamus, or a general feature of subcortical structures? To test the specificity of these thalamic moderation effects, we examined the pons, a brainstem structure within the hindbrain that is in the descending path from cerebral cortex to the cerebellar cortex, and not in the ascending pathway (**Fig 2c**). The cerebral cortex provides descending excitatory feedback to the cerebellar cortex via the pontine nuclei. However, we found a negative relationship between sensorimotor cortex and pons (β=−40.3, se=13.92, Cohen’s *f*=0.15, *p*=0.004). Pontine nuclei send signals to the cerebellar cortex via excitatory projections, and we found positive relationships between pons and cerebellar cortex (β=327.8, se=50.4, Cohen’s *f*=0.34, *p*<0.0001). Relationships did not differ between diagnosis groups in models controlling for sex, age, and eTIV. In a model that used the pons as a moderating variable across age groups (cerebellar cortex ~ SMC x age x pons) we found no significant effect of pons x age interaction (**Figure 2d-e**) [**Table S3**]. Thus, a difference in moderation effects between TD and ASD groups was found for thalamus, but not for pons. This finding is consistent with the concept that moderation effects arise from ascending signals in which cerebellum influences growth processes via thalamocortical pathways.

### Nonlinear Cognitive Trajectories Converge at Onset of Adolescence

The age-dependent decrease in thalamic moderation of cerebellum-neocortical covariation led us to examine whether changes in behavior followed a similar trajectory. Participating families completed the *BRIEF* questionnaire whose subsections assessed behavior in six separate cognitive domains, with higher scores indicating greater impairment (**Figure 3**). For each domain, we examined the trajectory of behavioral development between groups through an interaction model (age^2^ x group, controlled for sex). All six models showed greater impairment in the ASD group than in the TD group. When comparing the trajectory of behavioral development between groups, a striking pattern emerged. In three of the behavioral measures, differences significantly decreased until pre-adolescence and then rose again: Inhibition (Cohen’s *f*=0.16, *p*=0.031) (**Figure 3a**), Planning (Cohen’s *f*=0.19, *p*=0.015) (**Figure 3b**), and Emotional Regulation (Cohen’s *f*=0.20, *p*=0.009) (**Figure 3c**). The other three measures showed similar trajectories but differences in their age-dependence between groups was not significant: Shifting (*p*=0.53), Organization (*p*=0.81), Working Memory (*p*=0.10), and Initiation scores (*p*=0.066). Combining all behavioral measures into a single composite impairment score showed a broad inverse-parabolic effect of significant differences between groups (Cohen’s *f*=0.17, *p*=0.022) (**Fig 3d**).

**Fig. 3.**
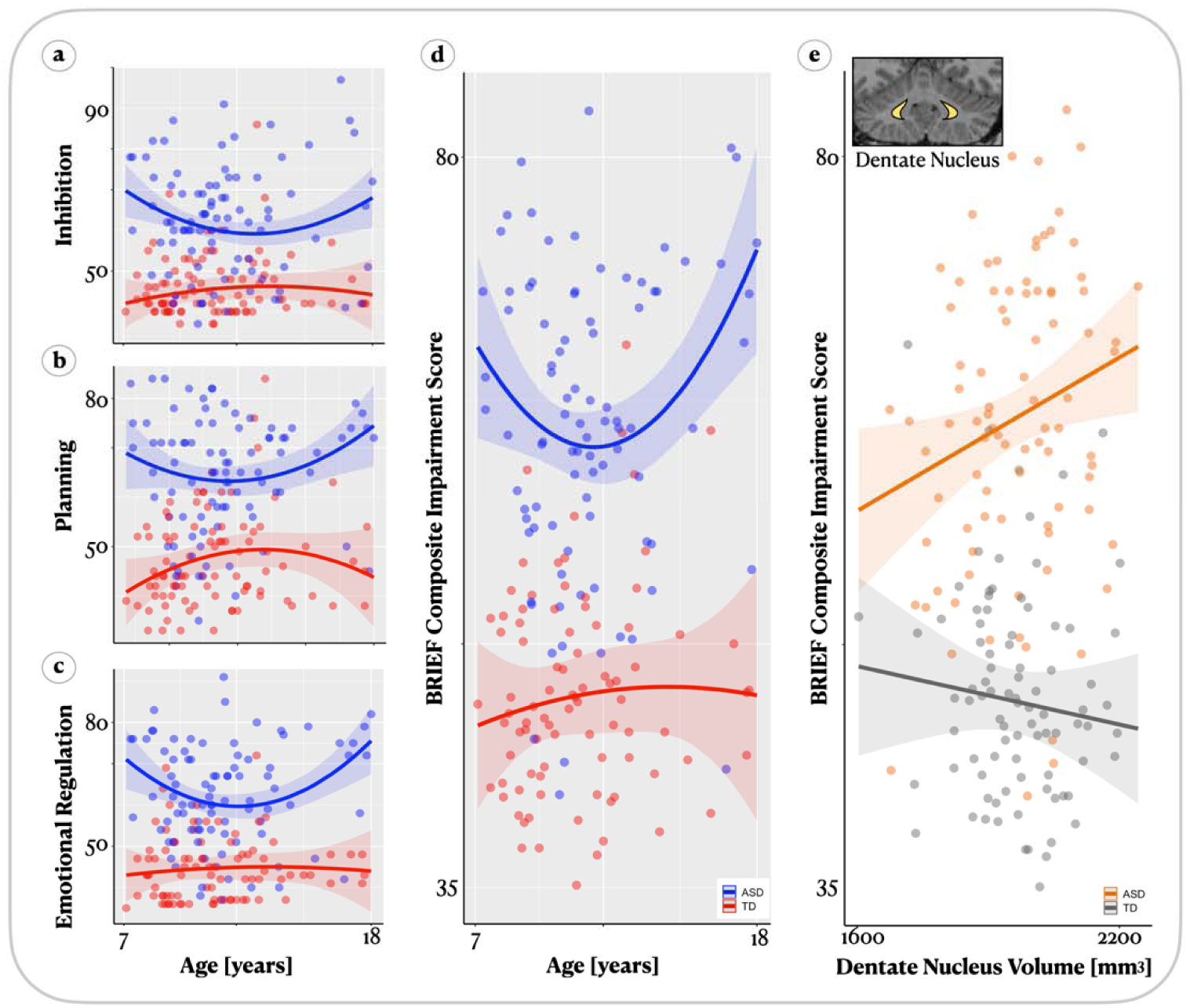
Cognitive trajectories converge in pre-adolescence and relate to cerebellum. Dimensions of cognitive function measured by the BRIEF parent-teacher questionnaire in a subset of individuals with complete behavioral data. In fitted models of age2 x group across sub­dimensions of cognitive function measured, corrected for sex, there was greater impairment in the autism cohort along (**a**) inhibition (β=−100.10, se=46.14, *p*=0.031), (**b**) planning (β=−119.62, se=48.69, *p*=0.015), and (**c**) emotional regulation (β=−117.55, se=44.85, *p*=0.009). Groups did not differ in their shifting (*p* = 0.790) or organization domains (*p*=0.786) (not shown). No cubic models were significant. When averaging all sub-scales into a composite measure (**d**), we found generally anti-correlated nonlinear cognitive trajectories between groups (β=−80.03, se=34.89, *p*=0.022). In ASD this was observed as a drop in impairment scores followed by a gradual rise with increasing age. (**e**) Further, correcting for biological sex, greater dentate cerebellar nuclei volume predicted greater impairment scores in ASD (β=−93.88, se=43.15, *p*=0.030).

Might the distinct behavioral trajectories between TD and ASD children result from neuroanatomical differences of the cerebello-thalamo-cortical network? We answered this question by correlating BRIEF cognitive scores with anatomy along the ascending cerebello-thalamo-SMC pathway. We focus on the dentate nucleus given its role as the source of ascending cerebellar signals in this tripartite network. In post-hoc tests controlling for age^2^, sex, and eTIV, we found that behavioral scores were related to dentate nuclei volumes differently in ASD and TD groups (Cohen’s *f*=0.16, *p*=0.030), with a stronger relationship in the ASD group. This effect was specific to the left dentate nucleus, the same region that showed different patterns of tripartite covariance with thalamus and sensorimotor cortex (**Figure 3e**). This effect was present even while dentate nuclei volumes did not differ between ASD and TD groups (controlling for sex, age, and eTIV: *p*=0.345).

### Whole-Neocortex Surface-Based Associations with Cerebellum

The previous results show that the thalamus differentially moderates the structural co-development of the cerebellum and neocortex in neurotypical and ASD children, with accompanying behavioral correlates. To more widely map the spatial distribution of this cerebellar influence, we derived maps of neocortical thickness for each brain, and calculated the correlation at each vertex with dentat nuclear volumes in a linear model that included sex and eTIV as covariates (neocortical thickness ~ sex + eTIV + DN volume x diagnosis), corrected for multiple comparisons (**Figure 4a**). ASD children displayed more correlated vertices than TD children, with associations found in bilateral posterior cingulate cortex, frontal and parietal operculum, lateral prefrontal cortex, auditory primary and association cortex, and lateral and inferior temporal cortex, including the lingual and ventral temporal cortex (**Figure 4a**).

**Figure 4.**
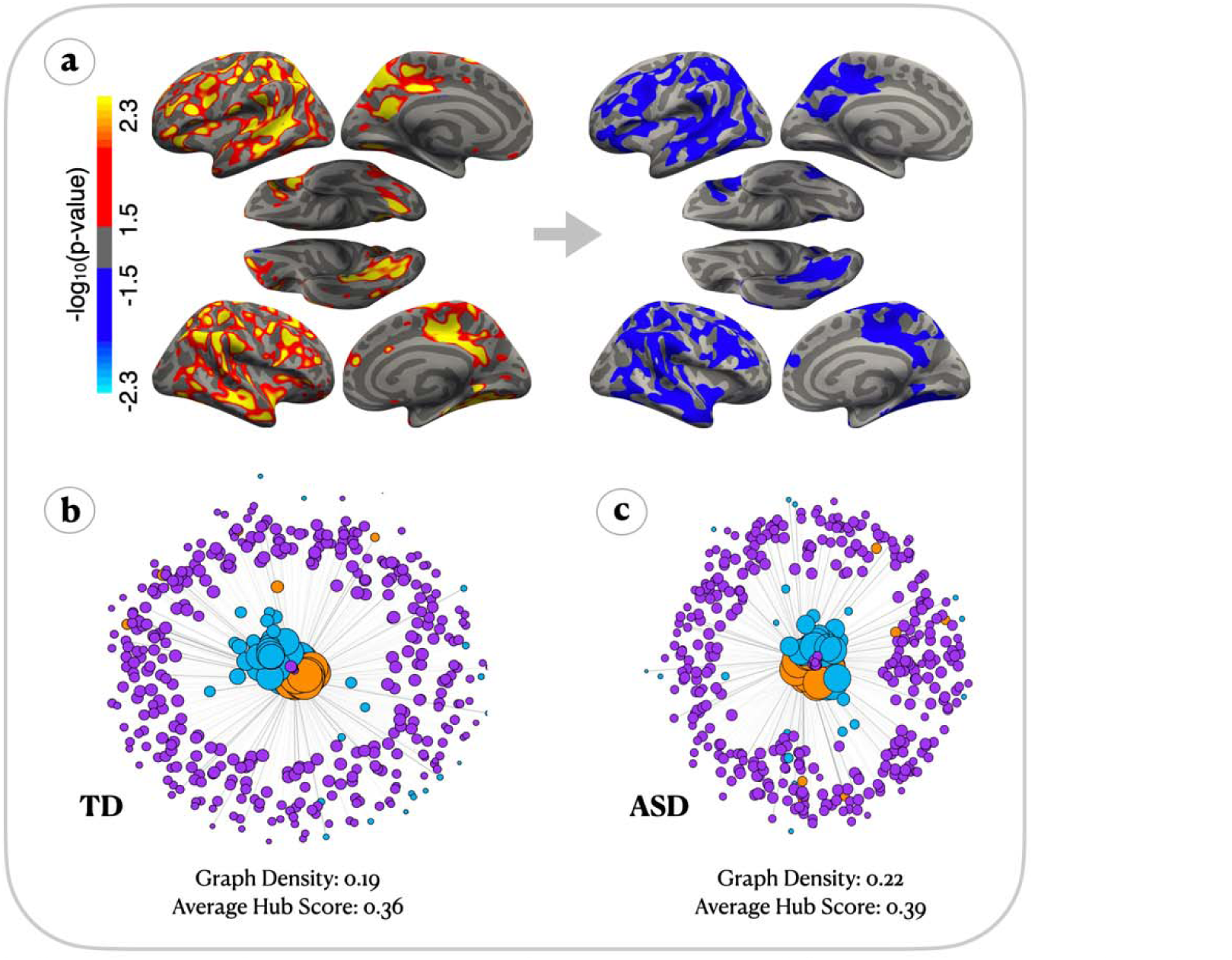
Cerebellar-Neocortical Covariance Network. (**a**) Vertex-wise differences in dentate nuclei-related cortical thickness, corrected for sex and intracranial volume, between autism and typically developing cohorts (*p*<0.05). Results surviving multiple-comparisons correction (a:right) encompassed left insular cortex, ventral temporal cortex, bilateral lateral occipital cortex, post central cortex, left opercularis, pars orbitalis, precuneus, and bilateral supra-marginal gyri, (**b-c**), Kamada-Kawai force-directed graphs for typically developing (**b**), and autism cohorts (**c**). Size of node represents nodal hub score, edges represent covariation, and the position of the node in the graph depicts its level of covariance to the rest of the network, with highly influential nodes placed in the middle and less influential nodes placed in the periphery. Purple=Neocortex, Blue=Cerebellum, Orange=Thalamic Nuclei.

With volume at each region, we further asked how regions of the cerebellum, thalamus, and neocortex covary as a larger network, treating the covariation between a given pair of regions, or nodes, as an edge in an extended graph. Such a graph analysis allowed us to examine if ASD was associated with network properties across the brain beyond sensorimotor cortex. In doing so, we explored the spatial extent to which the cerebellum might atypically correlate with cortex in ASD. To accomplish this analysis, we parcellated the neocortex into n=180 regions using the HCP-MMP atlas, and incorporated it into an unthresholded matrix of thalamic nuclei and cerebellar regions. These relationships can be visualized as Kamada-Kawai force-directed graphs, emphasizing central nodes highly correlated with other regions and more peripheral structures that show less covariation with other structures (**Figure 4b-4c**). Thalamic nuclei and cerebellar tissue (blue and orange nodes) play a visible, central role with sparser coupling to neocortical regions (purple nodes) in TD (**Fig 4b**) compared to ASD (**Fig 4c**). We next formally quantified differences in network properties between TD and ASD children. As expected, the thalamic nuclei and cerebellar nodes were the nodes with the greatest hub scores, a finding in line with their widespread connectivity to neocortex. A comparison of average graph properties, including graph density and hub score, of the cerebello-thalamocortical network between TD and ASD revealed that in ASD, the average nodal hub score was significantly greater relative to TD [*t*(860)=3.28, *p*=0.001], suggesting that ASD cerebellar and thalamic nodes were more broadly correlated with downstream cortical regions. There were no significant differences for nodal degree (ASD mean: 78.31, TD mean: 68.04, *p*=0.101) despite ASD trending toward greater degree values, suggesting that in ASD, brain regions tend to covary more with other regions than in TD.

## DISCUSSION

We found that the brains of children with autism showed different patterns of covariation between brain regions compared to neurotypicals, with the thalamus more strongly moderating structural covariance between cerebellar dentate nuclei and sensorimotor neocortex. These differences were prominent in childhood and decreased by adolescence, a decrease that paralleled the convergence in behavioral maturation between the two groups. These results, in line with the cerebellar diaschisis hypothesis of ASD, suggest that in autism there is an atypical early developmental influence from the cerebellum on brain growth via thalamocortical pathways.

The developmental diaschisis hypothesis proposes that the cerebellum plays a crucial role in postnatal development by influencing the maturation of distant brain regions to which it projects^2^. Cerebellar hypoplasia (*i.e.*, smaller volume) in ASD has been reported as early as 6 months^42^ to three years of age^43^, and here we found smaller cerebellar volumes in a cohort of 7– 18-year-olds (average age, 12-13 years old). Our analysis demonstrates widespread effects of cerebellar alterations on the development of the cortex, and suggests a means by which cerebellar hypoplasia could account for cortical structural abnormalities in ASD: Compromised cerebellar influence over neocortical processes stemming from smaller cerebella recall previous reports of larger neocortical sheets^44^ and increased sheet thickness^45^. In line with these studies, we found larger sensorimotor cortex and smaller cerebellar cortex in ASD [**Table 1**].

The present data link disparate findings within the field – less cerebellar tissue and increased cortical tissue – through a network model in which structures of the cerebellum, thalamus, and cortex covary in their volume according to the sign of their neuronal connections. Previous reports have documented positive coupling in functional activity between cerebellar and cerebral cortex^46,47^. Whether this was from the all-excitatory cortico-pontocerebellar descending pathway, or the ascending pathway that is characterized in the present study, was not clear. Here, by comparing thalamic-moderated effects *vs* pons-moderated effects in a structural covariance framework, we offer evidence that it is cerebellar output that influences cerebral cortex during development via the ascending cerebello-thalamocortical pathway.

Conflicting findings of cerebellar volumetric differences in ASD have previously been reported. Reports of no cerebellar size differences^48–51^ had fewer subjects than the present study (13 to 51 ASD cases, per study). One report of increased cerebellar volume did not correct for head size^52^. Two reports showing increased cerebellar volume that did consider total intracranial volume^53,54^ used processing pipelines not designed specifically for segmenting cerebellar gray matter volumes^53^ and again had few subjects (22 ASD cases^54^). While other reports demonstrated reduced cerebellar size similar to the present results [**Table 1**], they included low numbers of subjects^43,55–58^. We analyzed large public repositories using recently developed automatic cerebellar segmentation pipelines. Large datasets allowed us to model more complex relationships such as thalamic moderation and quadratic age effects. We further corrected for total head size and other confounding variables. To our knowledge, our approach is the first to account for these considerations in the assessment of cerebellar volumetry in ASD.

Neuroimaging and postmortem studies in ASD have previously shown that cerebellar abnormalities such as Purkinje cell loss in early life persist into adulthood^59,60^. In the present study we find that thalamic structural properties atypically influence correlations between the cerebellum and sensorimotor cortex in ASD. The inverse covariation between cerebellar cortex and thalamic/sensorimotor neocortical regions is consistent with cerebellar inhibition of thalamocortical excitatory systems^61^. While it has been proposed that the cerebellum is involved in ASD and its associated behavioral differences^62^, our findings extend this theory to encompass the idea that autism-related atypicality in cerebellar tissue also affects the structural development of long-range cortical networks. Furthermore, we provide evidence of the behavioral relevance of this structural variation. We find that greater volume of the cerebellar dentate nucleus relates to greater overall cognitive impairment in ASD, while for neurotypical individuals the slope of the relationship is flat or inverted, suggesting that dentate morphometry has different long-distance effects depending on diagnosis. The atypical lateralization we find also calls to mind previous reports of motor-system asymmetries in ASD^63–65^. Additionally, this asymmetry may reflect developmental neural shifts, such as those observed for implicit reading tasks involving a decrease in functional connectivity of the right cerebellar hemisphere in adult controls (older than 20 years) relative to control children (younger than 9 years)^66^, along with other widespread changes in the organizational architecture of functional brain networks believed to derive from age-dependent changes in gray matter^67^.

Previous observations suggest that in ASD, neocortical brain overgrowth differences which are observed in early childhood^44^ may diminish by adolescence^45^. This phenomenon appears to parallel a pre-adolescent ‘slump’ phase of cognitive development, a critical period during which the brain undergoes significant structural changes^68^. The term adolescent ‘slump’ refers to an observed temporary decline in academic performance during early adolescence, characterized by challenges in schoolwork, compared to earlier childhood or later adolescence^69,70^. We found that behavioral differences between ASD and non-ASD children diminished during early adolescence along with the strength of structural coupling between cerebellum, thalamus, and neocortex. These observations suggest that early developmental influences of cerebellar output on brain growth in ASD may be followed by further, compensatory change later in childhood^71^. We find such an instance in our dataset, in that ASD individuals experienced an improvement in their behavioral symptoms at pre-adolescent stages. This emphasizes the importance of considering the developmental trajectory of ASD, as studying adults alone may overlook these critical periods of change and hinder a comprehensive understanding of the neurobiological underpinnings of the disorder.

One potential avenue of future research is large-scale longitudinal investigation of biological mechanisms underlying the development of the cerebellum and neocortex within ASD and neurotypical individuals. A recent genome-wide association study (GWAS) showed a heritability (h^2^) of 0.51 for cerebellar volume, with enrichment for ASD-associated loci^72^. That study found 21 unique genes of interest for follow-up, as well as genetic correlations between total cerebellar volume and thalamic volume. Development-related shifts in gene-brain network associations may also occur. For example, childhood-to-adolescence shifts in network coupling patterns^73,74^ may stem from synaptic and myelin-related processes that peak in adolescence^75,76^. Recent work charting the fine-scale tissue development within the typical cerebellum^77^ invites investigation into myelin and other macromolecule expression in ASD. These findings suggest the possibility of identifying specific causes for the covariation we have observed.

### Conclusions

This study showed that children with autism have different patterns of coordinated brain development compared to neurotypical individuals, involving the cerebellum, thalamus, and sensorimotor cortex. These differences were larger in childhood and diminished in adolescence. These findings are consistent with the possibility that the cerebellum influences brain development via long-distance projections. The developmental approach employed here provides a baseline for studying different stages of development and for enhancing our understanding of pre-adolescent childhood as a key period of structural and behavioral change in ASD.

## Supplementary material

Supplementary material is found below.

## Abbreviations

ABIDE = Autism Brain Imaging Data Exchange; ASD = autism spectrum disorder; BRIEF = Behavior Rating Inventory of Executive Function; DN=dentate nuclei; TD = typically developing

---

**Figure S1.**
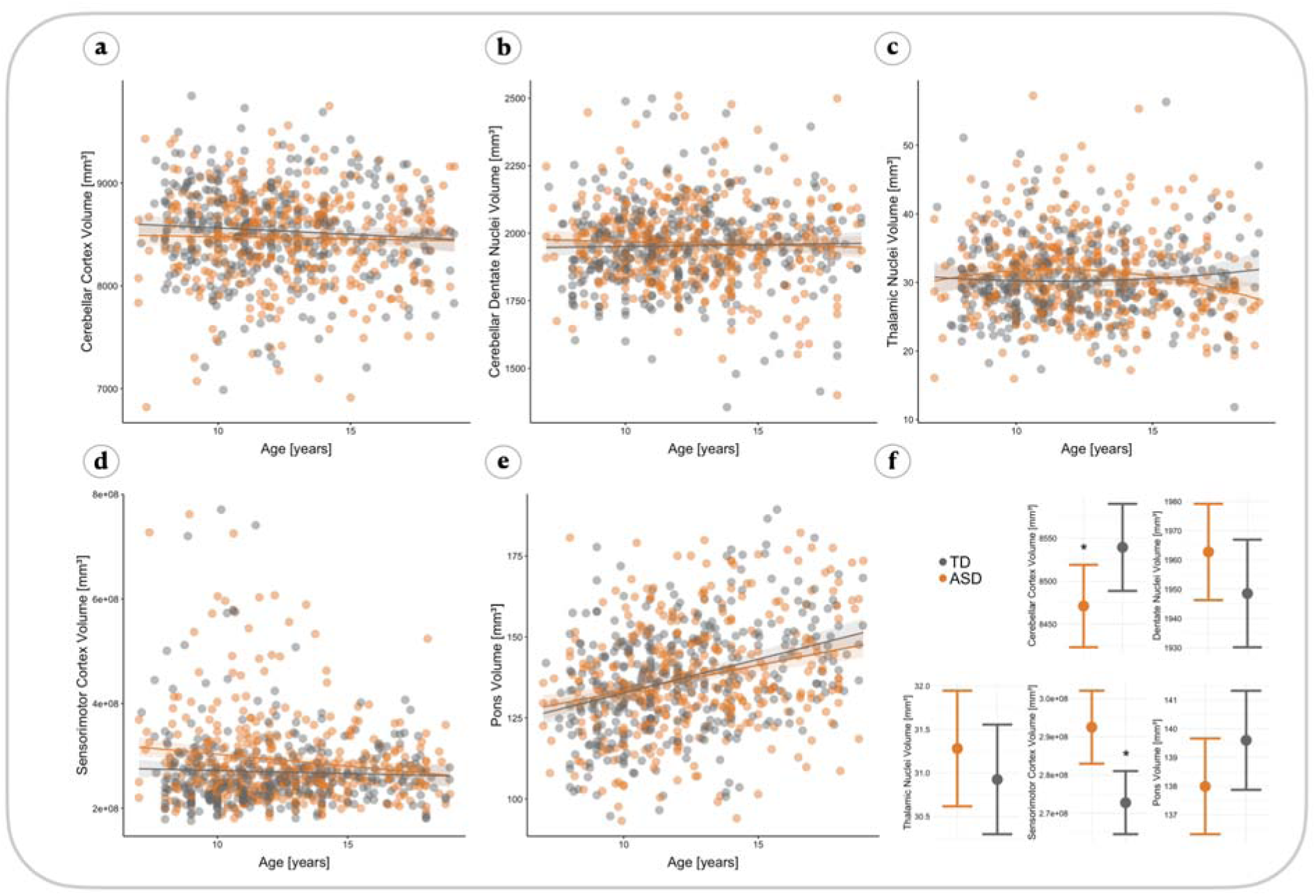
Age trajectories for main analyses ROIs. Average curve plots of relationships between age and gray matter volumes from cerebellar cortex (**a**), dentate cerebellar nuclei (**b**), thalamic nuclei (**c**), sensorimotor cortex (**d**), and pons (e). Correcting for sex and intracranial volume, as seen in f, group volumes differed on average in cerebellar cortex (TD mean: 8536.76, ASD mean: 8466.50; t(745)=2.06, ci=−[3.30,133.47], *p*=0.040) (**a**), and sensorimotor cortex (TD mean: 2696.48, ASD mean: 2919.76; t(745)=−3.33, ci=[−320.88, −83.26], *p*=0.001) (**d**). No significant group differences in average volume were found for thalamic nuclei (*p*=0.412), dentate cerebellar nuclei (p=0.225), or pons (p=0.159). Correcting for sex and intracranial volume, groups did not differ in their age trajectories (i.e., age x group effects) for cerebellar cortex volume *p*=0.485) (a), dentate cerebellar nuclei (*p*=0.292) (**b**), thalamic nuclei (β=0.26, se=0.15, *p*=0.090) (c), sensorimotor cortex (p=0.157) (d), or pons (p=0.641) (e).

**Table S1a.**
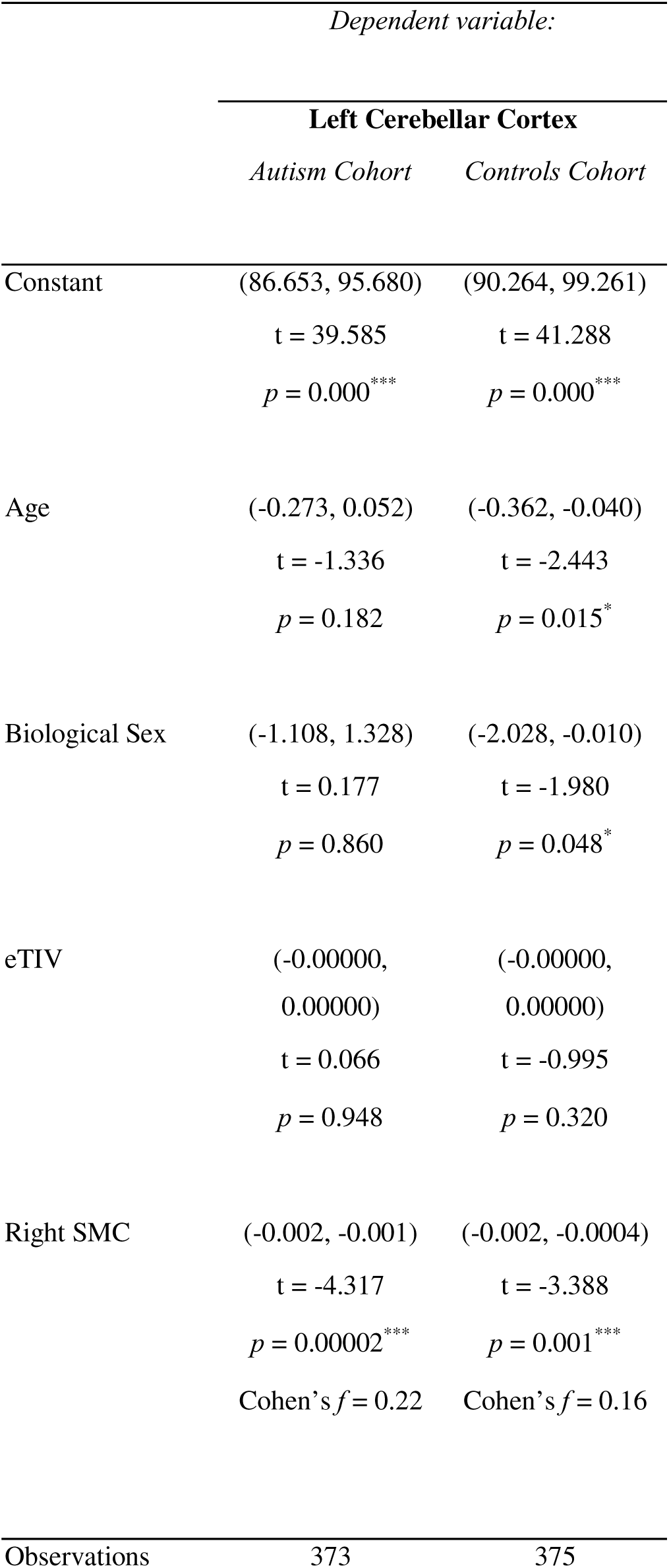
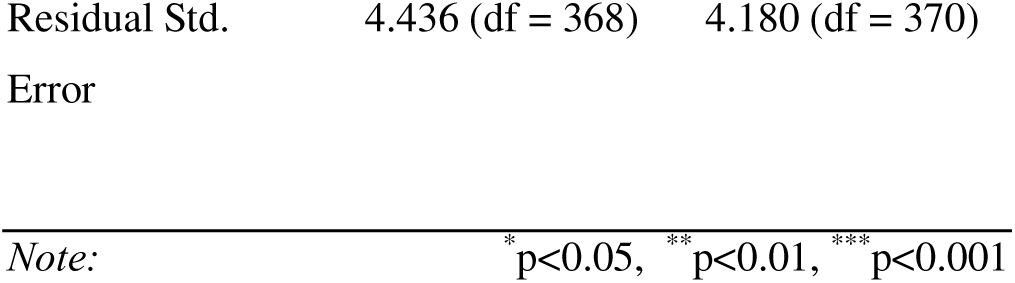

**Table S1b.**
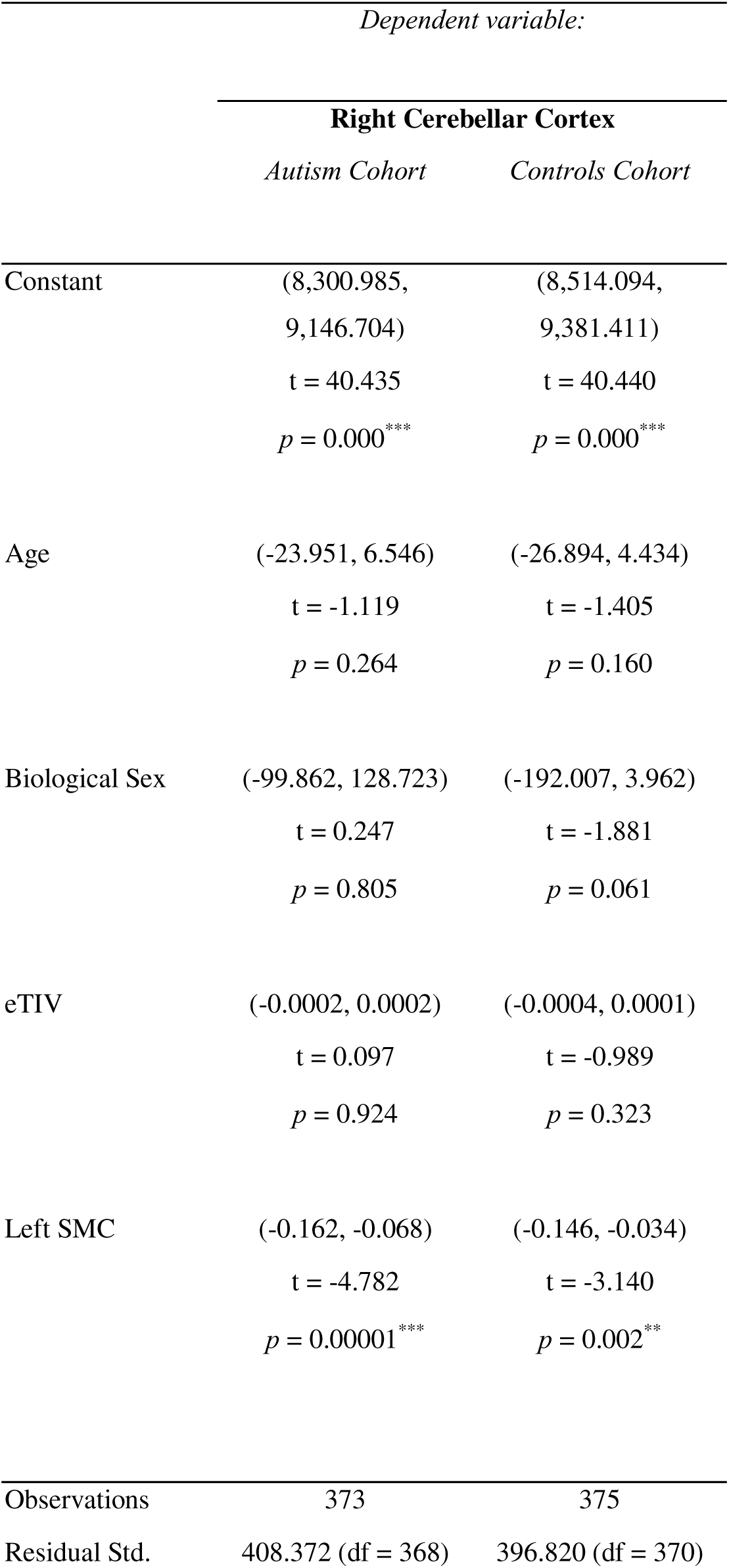
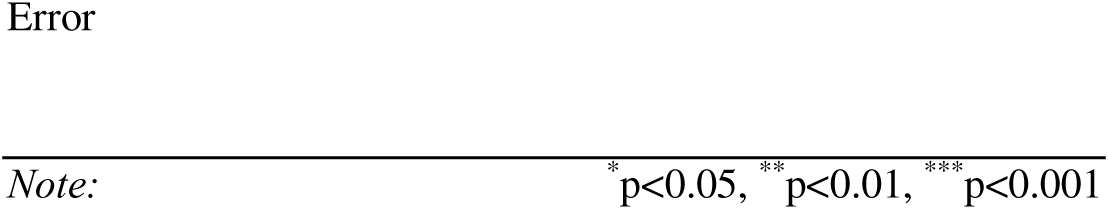

**Table S2.**
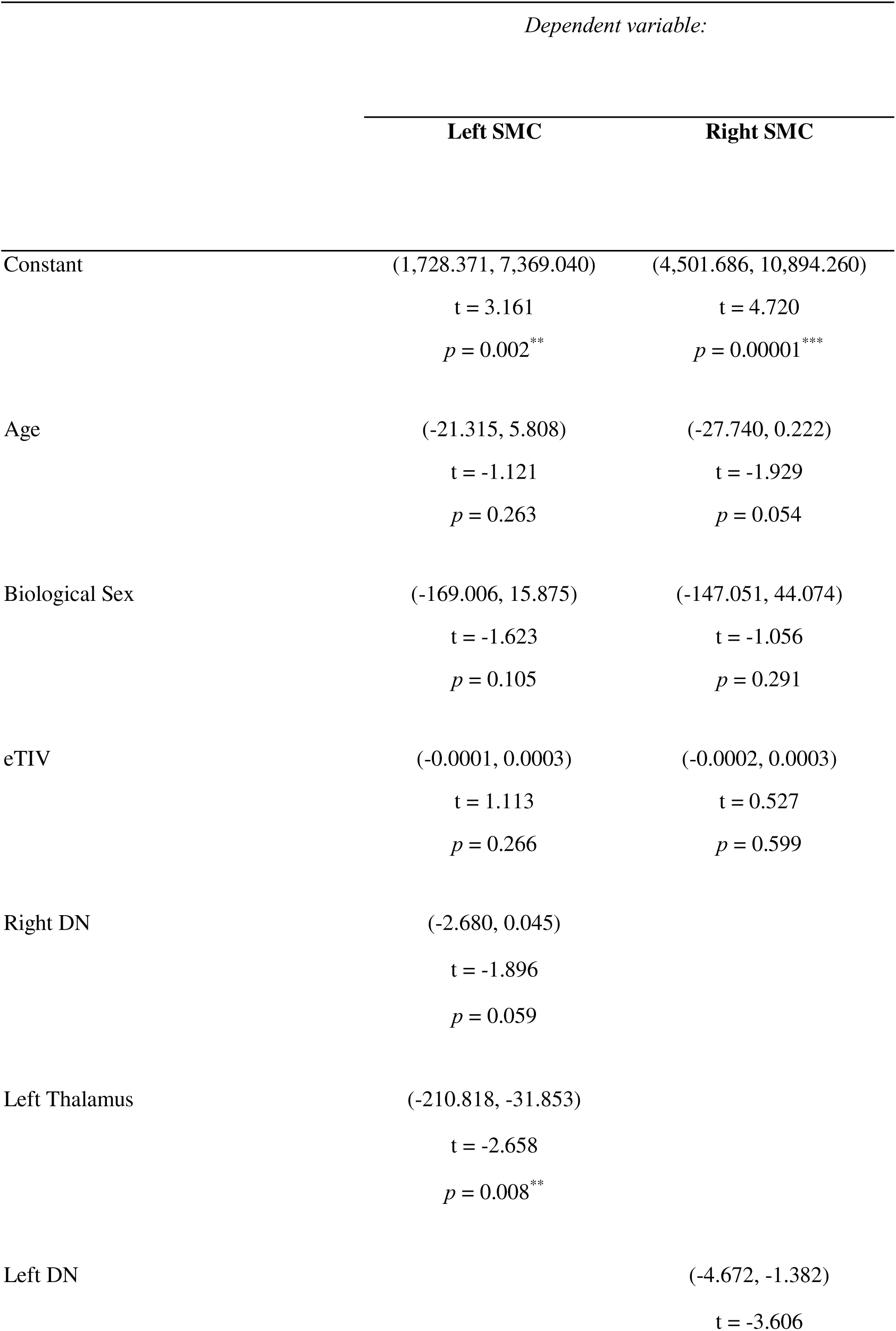
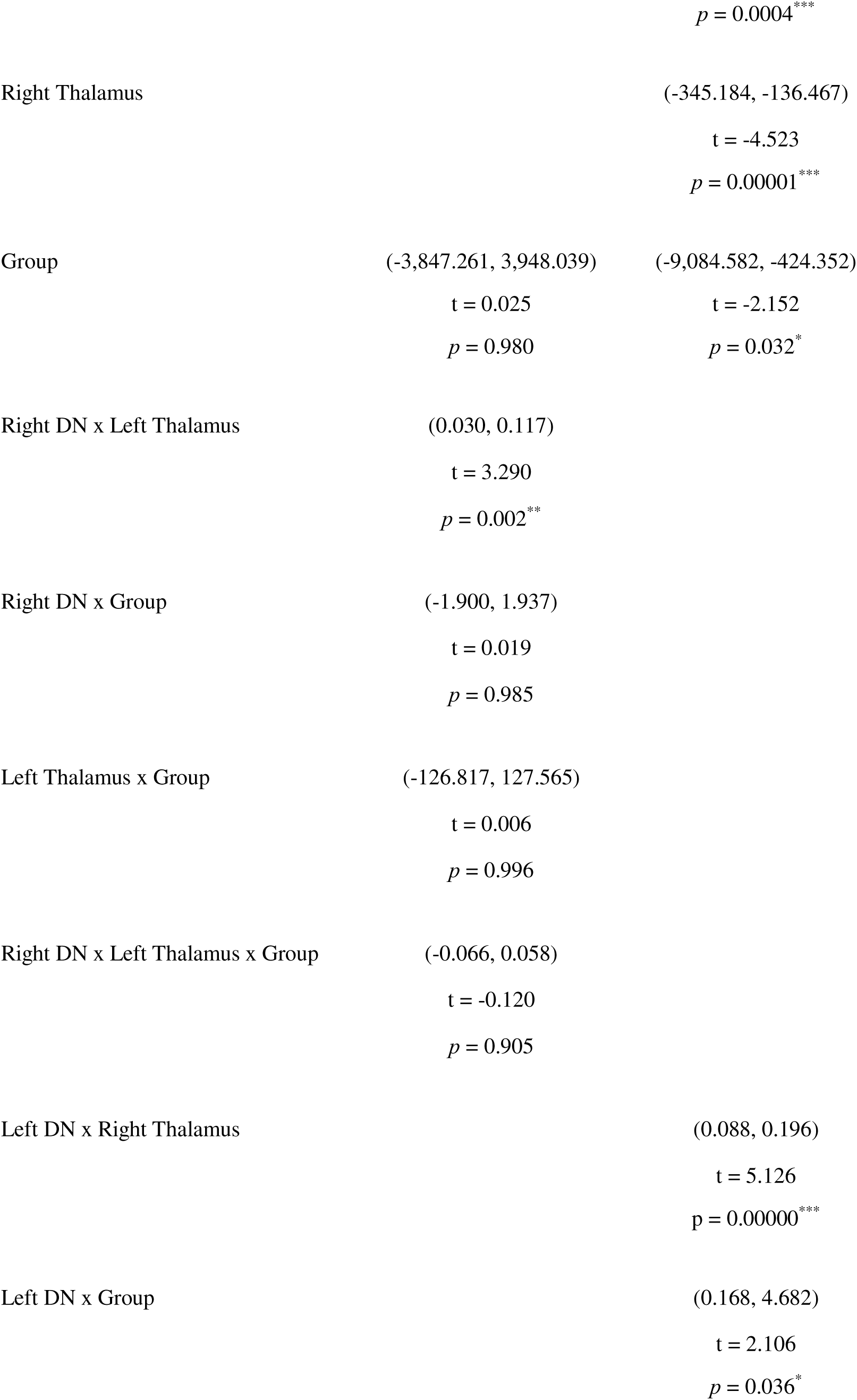
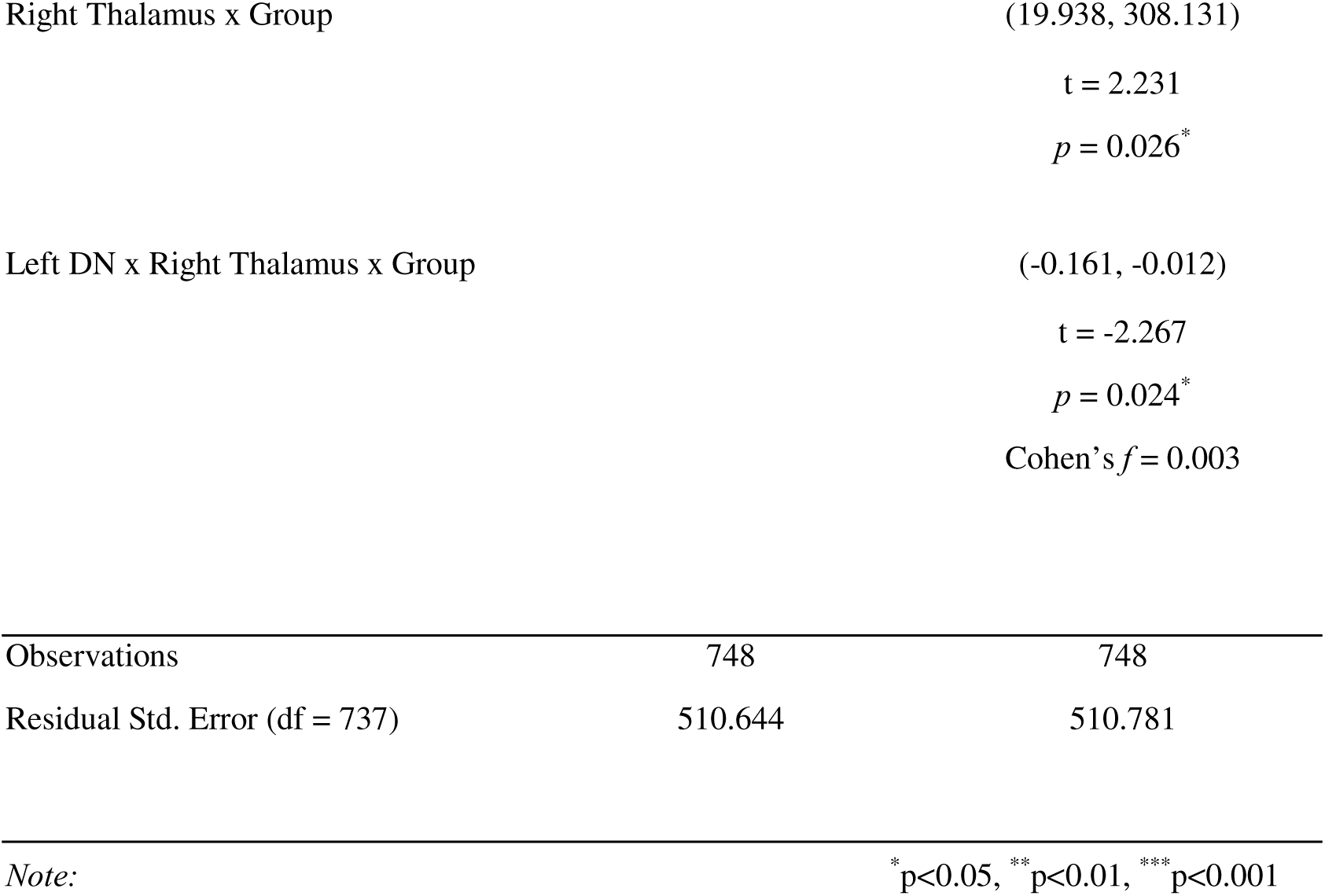

**Table S3a.**

| *Dependent variable:* |  |  |
| --- | --- | --- |
|  | **Left Cerebellar Cortex** |  |
|  | *Autism Cohort* | *Controls Cohort* |
| Constant | (7,741.221,<br>19,718.470)<br>t = 4.494<br>p = 0.00001*** | (3,504.908,<br>17,190.360)<br>t = 2.964<br>p = 0.004** |
| Right SMC | (-3.581, 0.597)<br>t = -1.400<br>p = 0.162 | (-3.038, 2.040)<br>t = -0.385<br>p = 0.701 |
| Pons | (-7.407, 1.498)<br>t = -1.301<br>p = 0.194 | (-5.144, 4.693)<br>t = -0.090<br>p = 0.929 |
| Age | (-937.437, -13.016)<br>t = -2.015<br>p = 0.044* | (-775.532, 408.650)<br>t = -0.607<br>p = 0.544 |
| Biological Sex | (-41.425, 186.692)<br>t = 1.248<br>p = 0.212 | (-146.147, 55.766)<br>t = -0.877<br>p = 0.381 |
| eTIV | (-0.001, -0.0001)<br>t = -2.725<br>p = 0.007** | (-0.001, -0.0002)<br>t = -3.168<br>p = 0.002** |
| Right SMC x Pons | (-0.001, 0.003) | (-0.002, 0.002) |
|  | t = 1.313 | t = 0.324 |
|  | p = 0.190 | p = 0.746 |
| Right SMC x Age | (-0.039, 0.282) | (-0.179, 0.263) |
|  | t = 1.480 | t = 0.377 |
|  | p = 0.139 | p = 0.707 |
| Pons x Age | (-0.010, 0.667) | (-0.314, 0.526) |
|  | t = 1.902 | t = 0.494 |
|  | p = 0.058 | p = 0.622 |
| Right SMC x Pons x Age | (-0.0002, 0.00003) | (-0.0002, 0.0001) |
|  | t = -1.471 | t = -0.378 |
|  | p = 0.142 | p = 0.706 |
| Observations | 374 | 375 |
| Residual Std. Error | 386.646 (df = 364) | 404.407 (df = 365) |
| *Note:* * p<0.05, ** p<0.01, *** p<0.001 |  |  |

**Table S3b.**

| *Dependent variable:* |  |  |
| --- | --- | --- |
| **Right Cerebellar Cortex** |  |  |
|  | *Autism Cohort* | *Controls Cohort* |
| Constant | (3,689.206,<br>14,831.020)<br>t = 3.258<br>p = 0.002** | (-835.811,<br>13,292.170)<br>t = 1.728<br>p = 0.084 |
| Left SMC | (-1.831, 1.853)<br>t = 0.012<br>p = 0.991 | (-1.760, 3.327)<br>t = 0.604<br>p = 0.546 |
| Pons | (-4.194, 4.108)<br>t = -0.020<br>p = 0.984 | (-2.800, 7.328)<br>t = 0.876<br>p = 0.381 |
| Age | (-595.615, 282.797)<br>t = -0.698<br>p = 0.486 | (-442.442, 782.232)<br>t = 0.544<br>p = 0.587 |
| Biological Sex | (-45.347, 173.686)<br>t = 1.148<br>p = 0.251 | (-137.522, 59.008)<br>t = -0.783<br>p = 0.434 |
| eTIV | (-0.001, -0.0001)<br>t = -2.830<br>p = 0.005** | (-0.001, -0.0002)<br>t = -3.288<br>p = 0.002** |
| Left SMC x Pons | (-0.001, 0.001) | (-0.002, 0.001) |
|  | t = -0.065 | t = -0.612 |
|  | p = 0.949 | p = 0.541 |
| Left SMC x Age | (-0.137, 0.153) | (-0.302, 0.142) |
|  | t = 0.107 | t = -0.708 |
|  | p = 0.915 | p = 0.479 |
| Pons x Age | (-0.219, 0.428) | (-0.567, 0.302) |
|  | t = 0.633 | t = -0.597 |
|  | p = 0.527 | p = 0.551 |
| Left SMC x Pons x Age | (-0.0001, 0.0001) | (-0.0001, 0.0002) |
|  | t = -0.144 | t = 0.677 |
|  | p = 0.886 | p = 0.499 |
| Observations | 374 | 375 |
| Residual Std. Error | 369.057 (df = 364) | 394.321 (df = 365) |

*Note:* \*p<0.05, \*\*p<0.01, \*\*\*p<0.001

## References

1. Schmahmann, J. D. The cerebellar cognitive affective syndrome: clinical correlations of the dysmetria of thought hypothesis. Int. Rev. Psychiatry (2001).

2. Wang, S. S.-H., Kloth, A. D. & Badura, A. The cerebellum, sensitive periods, and autism. Neuron 83, 518–532 (2014).

3. Limperopoulos, C. et al. Injury to the premature cerebellum: outcome is related to remote cortical development. Cereb. Cortex 24, 728–736 (2014).

4. Limperopoulos, C., Chilingaryan, G., Guizard, N., Robertson, R. L. & Du Plessis, A. J. Cerebellar injury in the premature infant is associated with impaired growth of specific cerebral regions. Pediatr. Res. 68, 145–150 (2010).

5. Zayek, M. M. et al. Cerebellar hemorrhage: a major morbidity in extremely preterm infants. J. Perinatol. 32, 699–704 (2012).

6. Steggerda, S. J. et al. Cerebellar Injury in Preterm Infants: Incidence and Findings on US and MR Images. Radiology 252, 190–199 (2009).

7. Newberg, A. B., Alavi, A. & Alavi, J. Contralateral cortical diaschisis in a patient with cerebellar astrocytoma after radiation therapy. Clin. Nucl. Med. 25, 431–433 (2000).

8. Sönmezoğlu, K., Sperling, B., Henriksen, T., Tfelt-Hansen, P. & Lassen, N. A. Reduced contralateral hemispheric flow measured by SPECT in cerebellar lesions: crossed cerebral diaschisis. Acta Neurol. Scand. 87, 275–280 (1993).

9. Rousseaux, M. & Steinling, M. Crossed hemispheric diaschisis in unilateral cerebellar lesions. Stroke 23, 511–514 (1992).

10. Botez, M. I., Léveillé, J., Lambert, R. & Botez, T. Single photon emission computed tomography (SPECT) in cerebellar disease: cerebello-cerebral diaschisis. Eur. Neurol. 31, 405–412 (1991).

11. Broich, K., Hartmann, A., Biersack, H. J. & Horn, R. Crossed cerebello-cerebral diaschisis in a patient with cerebellar infarction. Neurosci. Lett. 83, 7–12 (1987).

12. Di Lazzaro, V. et al. Excitability of the motor cortex to magnetic stimulation in patients with cerebellar lesions. J. Neurol. Neurosurg. Psychiatry 57, 108–110 (1994).

13. Moberget, T. et al. Long-term supratentorial brain structure and cognitive function following cerebellar tumour resections in childhood. Neuropsychologia 69, 218–231 (2015).

14. Cianfoni, A. et al. MRI findings of crossed cerebellar diaschisis in a case of Rasmussen’s encephalitis. J. Neurol. 257, 1748–1750 (2010).

15. Tien, R. D. & Ashdown, B. C. Crossed cerebellar diaschisis and crossed cerebellar atrophy: correlation of MR findings, clinical symptoms, and supratentorial diseases in 26 patients. AJR Am. J. Roentgenol. 158, 1155–1159 (1992).

16. Bond, K. M. et al. Dentate Update: Imaging Features of Entities That Affect the Dentate Nucleus. AJNR Am. J. Neuroradiol. 38, 1467–1474 (2017).

17. Asanuma, C., Thach, W. R. & Jones, E. G. Anatomical evidence for segregated focal groupings of efferent cells and their terminal ramifications in the cerebellothalamic pathway of the monkey. Brain Res. 286, 267–297 (1983).

18. Middleton, F. A. & Strick, P. L. Cerebellar projections to the prefrontal cortex of the primate. J. Neurosci. 21, 700–712 (2001).

19. Dum, R. P. & Strick, P. L. An unfolded map of the cerebellar dentate nucleus and its projections to the cerebral cortex. J. Neurophysiol. 89, 634–639 (2003).

20. Pisano, T. J. et al. Homologous organization of cerebellar pathways to sensory, motor, and associative forebrain. Cell Rep. 36, 109721 (2021).

21. Marco, E. J. et al. Children with autism show reduced somatosensory response: an MEG study. Autism Res. 5, 340–351 (2012).

22. Khan, S. et al. Somatosensory cortex functional connectivity abnormalities in autism show opposite trends, depending on direction and spatial scale. Brain 138, 1394–1409 (2015).

23. Alexander-Bloch, A., Giedd, J. N. & Bullmore, E. Imaging structural co-variance between human brain regions. Nat. Rev. Neurosci. 14, 322–336 (2013).

24. Zielinski, B. A., Gennatas, E. D., Zhou, J. & Seeley, W. W. Network-level structural covariance in the developing brain. Proc. Natl. Acad. Sci. U. S. A. 107, 18191–18196 (2010).

25. Di Martino, A. et al. Enhancing studies of the connectome in autism using the autism brain imaging data exchange II. Sci Data 4, 170010 (2017).

26. Di Martino, A. et al. The autism brain imaging data exchange: towards a large-scale evaluation of the intrinsic brain architecture in autism. Mol. Psychiatry 19, 659–667 (2014).

27. Lord, C., Rutter, M. & Le Couteur, A. Autism Diagnostic Interview-Revised: a revised version of a diagnostic interview for caregivers of individuals with possible pervasive developmental disorders. J. Autism Dev. Disord. 24, 659–685 (1994).

28. Michael; Lecouteur Rutter (Ann; Lord & Catherine). *Autism diagnostic interview: revised (ADI-R)*. (Western Psychological Services.).

29. Lord, C., Rutter, M., DiLavore, P. C. & Risi, S. Autism Diagnostic Observation Schedule: Ados-2. (Western Psychological Services, 2006).

30. Lord, C. et al. The Autism Diagnostic Observation Schedule—Generic: A Standard Measure of Social and Communication Deficits Associated with the Spectrum of Autism. J. Autism Dev. Disord. 30, 205–223 (2000).

31. Kaufman, J. & Schweder, A. E. The Schedule for Affective Disorders and Schizophrenia for School-Age Children: Present and Lifetime version (K-SADS-PL). in Comprehensive handbook of psychological assessment, Vol (ed. Hilsenroth, M. J.) vol. 2 247–255 (John Wiley & Sons, Inc., xvi, 2004).

32. First, M., Spitzer, R., Williams, J. & Gibbon, M. Structured clinical interview for DSM-IV—non-patient edition, version 1.0. Washington, DC: American Psychiatric.

33. Gioia, G. A., Isquith, P. K., Guy, S. C. & Kenworthy, L. Behavior rating inventory of executive function: BRIEF. (Psychological Assessment Resources Odessa, FL, 2000).

34. Fischl, B., Sereno, M. I. & Dale, A. M. Cortical surface-based analysis. II: Inflation, flattening, and a surface-based coordinate system. Neuroimage 9, 195–207 (1999).

35. Dale, A. M., Fischl, B. & Sereno, M. I. Cortical surface-based analysis. I. Segmentation and surface reconstruction. Neuroimage 9, 179–194 (1999).

36. Iglesias, J. E. et al. A probabilistic atlas of the human thalamic nuclei combining ex vivo MRI and histology. Neuroimage 183, 314–326 (2018).

37. Glasser, M. F. et al. A multi-modal parcellation of human cerebral cortex. Nature 536, 171–178 (2016).

38. Diedrichsen, J. A spatially unbiased atlas template of the human cerebellum. Neuroimage 33, 127–138 (2006).

39. Diedrichsen, J. et al. Imaging the deep cerebellar nuclei: a probabilistic atlas and normalization procedure. Neuroimage 54, 1786–1794 (2011).

40. Greve, D. N. & Fischl, B. False positive rates in surface-based anatomical analysis. Neuroimage 171, 6–14 (2018).

41. Anteraper, S. A. et al. Intrinsic Functional Connectivity of Dentate Nuclei in Autism Spectrum Disorder. Brain Connect. 9, 692–702 (2019).

42. Hashimoto, T. et al. Development of the brainstem and cerebellum in autistic patients. J. Autism Dev. Disord. 25, 1–18 (1995).

43. Courchesne, E. et al. Unusual brain growth patterns in early life in patients with autistic disorder: an MRI study. Neurology 57, 245–254 (2001).

44. Hazlett, H. C. et al. Early brain development in infants at high risk for autism spectrum disorder. Nature 542, 348–351 (2017).

45. Khundrakpam, B. S., Lewis, J. D., Kostopoulos, P., Carbonell, F. & Evans, A. C. Cortical Thickness Abnormalities in Autism Spectrum Disorders Through Late Childhood, Adolescence, and Adulthood: A Large-Scale MRI Study. Cereb. Cortex 27, 1721–1731 (2017).

46. Xue, A. et al. The detailed organization of the human cerebellum estimated by intrinsic functional connectivity within the individual. J. Neurophysiol. 125, 358–384 (2021).

47. Buckner, R. L., Krienen, F. M., Castellanos, A., Diaz, J. C. & Yeo, B. T. T. The organization of the human cerebellum estimated by intrinsic functional connectivity. J. Neurophysiol. 106, 2322–2345 (2011).

48. Hazlett, H. C. et al. Magnetic resonance imaging and head circumference study of brain size in autism: birth through age 2 years. Arch. Gen. Psychiatry 62, 1366–1376 (2005).

49. Courchesne, E., Townsend, J. & Saitoh, O. The brain in infantile autism: posterior fossa structures are abnormal. Neurology 44, 214–223 (1994).

50. Holttum, J. R., Minshew, N. J., Sanders, R. S. & Phillips, N. E. Magnetic resonance imaging of the posterior fossa in autism. Biol. Psychiatry 32, 1091–1101 (1992).

51. Manes, F. et al. An MRI study of the corpus callosum and cerebellum in mentally retarded autistic individuals. J. Neuropsychiatry Clin. Neurosci. 11, 470–474 (1999).

52. Piven, J., Saliba, K., Bailey, J. & Arndt, S. An MRI study of autism: the cerebellum revisited. Neurology 49, 546–551 (1997).

53. Traut, N. et al. Cerebellar Volume in Autism: Literature Meta-analysis and Analysis of the Autism Brain Imaging Data Exchange Cohort. Biol. Psychiatry 83, 579–588 (2018).

54. Hardan, A. Y., Minshew, N. J., Harenski, K. & Keshavan, M. S. Posterior fossa magnetic resonance imaging in autism. J. Am. Acad. Child Adolesc. Psychiatry 40, 666–672 (2001).

55. Hodge, S. M. et al. Cerebellum, language, and cognition in autism and specific language impairment. J. Autism Dev. Disord. 40, 300–316 (2010).

56. Scott, J. A., Schumann, C. M., Goodlin-Jones, B. L. & Amaral, D. G. A comprehensive volumetric analysis of the cerebellum in children and adolescents with autism spectrum disorder. Autism Res. 2, 246–257 (2009).

57. Piven, J. et al. Magnetic resonance imaging in autism: measurement of the cerebellum, pons, and fourth ventricle. Biol. Psychiatry 31, 491–504 (1992).

58. Courchesne, E., Yeung-Courchesne, R., Press, G. A., Hesselink, J. R. & Jernigan, T. L. Hypoplasia of cerebellar vermal lobules VI and VII in autism. N. Engl. J. Med. 318, 1349–1354 (1988).

59. Palmen, S. J. M. C., van Engeland, H., Hof, P. R. & Schmitz, C. Neuropathological findings in autism. Brain 127, 2572–2583 (2004).

60. Bailey, A. et al. A clinicopathological study of autism. Brain 121 (Pt 5), 889–905 (1998).

61. Heck, D. H., Fox, M. B., Correia Chapman, B., McAfee, S. S. & Liu, Y. Cerebellar control of thalamocortical circuits for cognitive function: A review of pathways and a proposed mechanism. Front. Syst. Neurosci. 17, 1126508 (2023).

62. Mapelli, L., Soda, T., D’Angelo, E. & Prestori, F. The Cerebellar Involvement in Autism Spectrum Disorders: From the Social Brain to Mouse Models. Int. J. Mol. Sci. 23, (2022).

63. Rysstad, A. L. & Pedersen, A. V. There Are Indeed More Left-Handers Within the Autism Spectrum Disorder Compared with in the General Population, but the Many Mixed-Handers Is the More Interesting Finding. J. Autism Dev. Disord. 48, 3253–3255 (2018).

64. Kobylinska, L., Anghel, C. G., Mihailescu, I., Rad, F. & Dobrescu, I. Handedness in Children with Autism Spectrum Disorders. Eur. Psychiatry 41, S214–S214 (2017).

65. Markou, P., Ahtam, B. & Papadatou-Pastou, M. Elevated Levels of Atypical Handedness in Autism: Meta-Analyses. Neuropsychol. Rev. 27, 258–283 (2017).

66. Turkeltaub, P. E., Gareau, L., Flowers, D. L., Zeffiro, T. A. & Eden, G. F. Development of neural mechanisms for reading. Nat. Neurosci. 6, 767–773 (2003).

67. Aboud, K. S. et al. Structural covariance across the lifespan: Brain development and aging through the lens of inter-network relationships. Hum. Brain Mapp. 40, 125–136 (2019).

68. Blakemore, S.-J., Burnett, S. & Dahl, R. E. The role of puberty in the developing adolescent brain. Hum. Brain Mapp. 31, 926–933 (2010).

69. Chall, J. S., Jacobs, V. A. & Baldwin, L. E. The Reading Crisis: Why Poor Children Fall Behind. (Harvard University Press, 1990).

70. Chall, J. S. Stages of Reading Development. (McGraw-Hill, 1983).

71. Livingston, L. A. & Happé, F. Conceptualising compensation in neurodevelopmental disorders: Reflections from autism spectrum disorder. Neurosci. Biobehav. Rev. 80, 729– 742 (2017).

72. Chambers, T. et al. Genetic common variants associated with cerebellar volume and their overlap with mental disorders: a study on 33,265 individuals from the UK-Biobank. Mol. Psychiatry 27, 2282–2290 (2022).

73. Váša, F. et al. Conservative and disruptive modes of adolescent change in human brain functional connectivity. Proc. Natl. Acad. Sci. U. S. A. 117, 3248–3253 (2020).

74. Schlesinger, K. J., Turner, B. O., Lopez, B. A., Miller, M. B. & Carlson, J. M. Age-dependent changes in task-based modular organization of the human brain. Neuroimage 146, 741–762 (2017).

75. Skene, N. G., Roy, M. & Grant, S. G. A genomic lifespan program that reorganises the young adult brain is targeted in schizophrenia. Elife 6, (2017).

76. Whitaker, K. J. et al. Adolescence is associated with genomically patterned consolidation of the hubs of the human brain connectome. Proc. Natl. Acad. Sci. U. S. A. 113, 9105–9110 (2016).

77. Liu, X., d’Oleire Uquillas, F., Viaene, A. N., Zhen, Z. & Gomez, J. A multifaceted gradient in human cerebellum of structural and functional development. Nat. Neurosci. 25, 1129–1133 (2022).

